# A Trypanosome Trifecta: an independently tunable Triple Inducer System for genetic studies in *Trypanosoma brucei*

**DOI:** 10.1101/2025.08.31.673413

**Authors:** Matt J. Romprey, Raveen Armstrong, Peter McGill, Michele M. Klingbeil

**Affiliations:** Department of Microbiology, University of Massachusetts, Amherst, MA, United States of America; Institute for Applied Life Sciences, University of Massachusetts, Amherst, MA, United States of America

## Abstract

*Trypanosoma brucei* is a model unicellular parasite for cellular and molecular genetic studies, but tools for more multiplexed experiments are limited. Tetracycline (Tet)-inducible gene regulation has been a long-standing and effective approach for overexpression, RNAi and genome wide screens. The single inducer Tet-On system has been foundational for studying essential genes that are required for biological processes and identifying potential drug targets. To achieve greater flexibility in experimental design, we capitalized upon previously described dual inducer systems that combined vanillic acid (Van) or cumate (Cym) with Tet as inducers. Here we report the development of a triple inducer system combining Cym, Van and Tet repressor elements to selectively regulate the expression of three genetic elements within a single cell line called PHITER. As proof of principle to demonstrate independent control, we adapted the previously characterized Van/Tet dual inducer RNAi complementation cell line, IBComp^VaT^ for additional Cym inducible expression of an eGFP reporter. We provide evidence that each of the inducer elements operate independently with no evidence of leaky expression. In this system, Cym induction is specific and tunable. Additionally, simultaneous triple induction for *POLIB* RNAi complementation with wildtype POLIB and eGFP expression phenocopied the near complete rescue of *POLIB* RNAi previously reported. This system enables the study of complex and pleiotropic phenotypes through more sophisticated experimental design.

## INTRODUCTION

The African trypanosome, *Trypanosoma brucei*, is an early diverging unicellular parasite that causes the Neglected Tropical Disease human African trypanosomiasis and a related wasting disease in cattle called nagana (1). In addition to medical and economic importance, *T. brucei* has served as a tractable model organism with many advanced genetics tools and insights into fundamental biological processes based on their distinctive biology. Examples include extensive post-transcriptional gene regulation, mitochondrial RNA editing which overturned years of molecular central dogma and unusual metabolic adaptations (2, 3). Another example of extreme biology is their mitochondrial DNA network called kinetoplast DNA (kDNA). Trypanosomes have one of the most topologically complex mitochondrial genomes in nature composed of maxicircles and minicircles that are catenated into a single network.

The kDNA structure and replication mechanism are divergent from all other eukaryotes and is essential for parasite survival and life cycle completion (4). A hallmark of kDNA replication is the topoisomerase II-mediated minicircle release and attachment mechanism, while maxicircles replicate still catenated within the network (5, 6, 7). Remarkably, minicircle progeny are attached to the network while still containing at least one gap. This “mark” on replicated progeny is a possible mechanism to ensure each circle is replicated just once per cell cycle. However, persistent gaps (ssDNA regions) negatively impact genome stability and can lead to replication stress, suggesting that the DNA damage tolerance response during kDNA replication is essential and therefore notably different in trypanosomes (8, 9).

The complexity of kDNA replication is also reflected in the many additional factors not present in mammalian nucleoids including 6 DNA polymerases (Pol), 6 helicases, 3 topoisomerases, 2 primases, 2 ligases and 2 origin binding proteins among others. Greater than 50 proteins have roles in the replication and segregation of the DNA network as well as maintaining the kDNA structural integrity and have been summarized elsewhere (10, 11). Interestingly, many of the kDNA replication factors have non-redundant roles discovered using RNAi for reverse genetic analysis of gene function. For example, *T. brucei* uses three family A DNA Pol paralogs (POLIB, POLIC, POLID) that are <u>independently essential</u> suggesting specialized roles in maintaining the kDNA network (12, 13, 14, 15). The 29-13 tetracycline (Tet)-On single inducer system has been crucial for single gene and genome-wide RNAi studies over the past 2 decades (16).

Among the many improvements to trypanosome molecular biology constructs, independent dual inducible systems were developed using Tet and vanillic acid (Van) or Tet and cumate (Cym) (17, 18). We recently reported the use of a dual inducer system for independently regulating RNAi and overexpression through the addition of Van and Tet, respectively. Simultaneous induction in this system allowed for robust RNAi complementation to study the kDNA replication protein POLIB (19). RNAi complementation studies and their associated control experiments can now be performed within a single cell line.

This highly tractable system could be expanded for further experimental flexibility by adding a third inducer, thus allowing for the independent regulation of 3 genetic elements in one cell line. In this triple inducer system, RNAi complementation could still operate using the Van and Tet inducer elements to concurrently control RNAi and overexpression, while the Cym inducer elements would allow for tunable regulation of an additional gene. A triple inducer system would allow expression of an additional gene during RNAi complementation, such as one that impacts kDNA genome stability. This can allow dissection of protein domain function during normal replication fork progression and possibly evaluate overlapping roles of kDNA replication and repair.

Here we describe the establishment of a triple inducer parental cell line that expresses all three repressor elements and demonstrate independent, and tunable Cym inducible expression using an eGFP reporter. To evaluate the full potential of a triple inducible system, we built upon the previously characterized Van/Tet dual inducer cell line for POLIB RNAi complementation (19) as proof of principle to demonstrate that all three elements can function independently and simultaneously. This tool overcomes key limitations and opens new avenues for trypanosomatid functional genomics.

## MATERIALS AND METHODS

For primer sequences refer to Table S1.

For cell lines and associated modifications used in this study refer to Table S2.

### DNA constructs

#### Single marker cumate inducible system

The hygromycin resistance gene was PCR amplified from pKOHyg (20) using primers UM147 and UM149 for subsequent Gibson assembly (NEB) with EcoRI and NcoI digested pSmOxNUS (18) to create pSmOxNUSHyg. The cumate repressor (CymR) gene and the hygromycin resistance gene (Hph) were then PCR amplified from pSmOxNUSHyg using primers UM159 and UM160 for Gibson assembly with StuI and BsiWI digested pDEX-CuO (18) to create pCuRO-eGFP.

#### Cumate repressor plasmid

The CymR and Hph genes were PCR amplified from pSmOxNUSHyg using primers UM192 and UM193 for subsequent Gibson assembly with NheI and SpeI digested pJ1173 (17) to create pCymRHyg thus removing the T7 RNA polymerase, tetracycline repressor (TetR) and a puromycin resistance gene from pJ1173.

#### Cumate inducible expression vector

The neomycin resistance gene was PCR amplified from pC-PTP-NEO (Shimanski 2005) using primers UM184 and UM185 for subsequent Gibson assembly (NEB) with BsiWI and StuI digested pDEX-CuO (18) to create pCuO-eGFP.

#### Trypanosome cell culture and transfection

The *Trypanosoma brucei brucei* procyclic cell line SMUMA (**S**ingle **M**arker **UMA**ss) contains the integrated plasmid pJ1173 (17) carrying T7 RNA polymerase (T7RNAP), tetracycline repressor (TetR) and vanillic acid repressor (VanR). SMUMA cells were cultured at 27°C in SDM-79 medium supplemented with 15% heat-inactivated fetal bovine serum and 1how to µg/ml puromycin (Puro) (19). The IBComp^VaT^ cell line (19) was cultured in SDM-79 medium supplemented with blasticidin (15 μg/mL), phleomycin (2.5 μg/mL) and puromycin (1 μg/mL).

To create the Cym triple inducer reporter cell line, CuRO-eGFP, NotI linearized pCuRO-eGFP was transfected into SMUMA cells via nucleofection to integrate into the 177 bp locus and selected with 50 μg/mL hygromycin (Hyg). To create the triple inducer PHITER parental cell line that expresses three repressor elements (CymR, TetR, VanR), SMUMA (19) was transfected with HindIII digested pCymRHyg via nucleofection to integrate into the α/β tubulin array and selected with 50 μg/mL Hyg. Verification of all three repressors elements in PHITER clones are provided in Fig S1. Four clonal cell lines were chosen based on doubling times and subsequently transfected via nucleofection with NotI linearized pCuO-eGFP for integration into the 177 bp repeat region and selected with 50 µg/mL G418 to screen for Cym inducible protein expression. These populations are called eGFP^PHITER^ and were not further dilution cloned. All other transfected cell lines were additionally supplemented with the appropriate selectable drug prior to limiting dilution to obtain clonal cell lines.

The POLIB dual inducer cell line IBComp^VaT^ clone P5D1 (19) was transfected with HindIII digested pCymRHyg and selected with 50 μg/mL Hyg. Clone P1D8 was chosen for this study based on doubling time and named IBComp^PHIT^ (<u>P</u>uro, <u>H</u>yg, Inducible <u>T</u>riad). IBComp^PHIT^ was subsequently transfected with NotI linearized pCuO-eGFP and selected with 50 µg/mL G418 to create the IBComp-eGFP^PHIT^ reporter cell line. Repressor expression was verified by northern blot and clone P1F12 was chosen for this study.

#### Inducible expression

Expression of eGFP was induced with 25 μg/mL of Cym for 48 hr for both triple inducer reporter cell lines CuRO-eGFP and eGFP^PHITER^. In the proof-of-concept cell line, IBComp-eGFP^PHIT^, RNAi was induced by addition of Van (250 µM dissolved in DMSO) and expression of an ectopic recoded version of wild type POLIB PTP-tagged variant (POLIBWT) was induced by addition of Tet (4 µg/mL). Cultures were supplemented daily with Van and/or Tet to maintain RNAi and protein expression (19). For eGFP expression, cells were induced with Cym (3 μg/mL dissolved in DMSO) on transfer days. To assess if Cym induction was reversible, IBComp-eGFP^PHIT^ cells were grown for 2 days in the presence of 3 μg/mL Cym only or in the presence of all three inducers. Cells were then pelleted, and then washed once in 1X phosphate buffered saline (PBS) to remove the inducer. Cells were resuspended and grown in media lacking Cym for an additional 3 days. Cells were then harvested at indicated time points for SDS-PAGE and western blot analyses.

#### SDS PAGE & western blotting

Cells were pelleted, washed with 1X PBS, supplemented with 1X protease inhibitor cocktail (Roche), resuspended and stored at −80°C. Protein samples were fractionated in a sodium-dodecyl-sulfate polyacrylamide gel (150 V) and transferred overnight (90 mA) onto a PVDF membrane. Membranes were then blocked in a 5% w/v non-fat dry milk solution for 2 hrs.

PTP-tagged POLIB was detected with peroxidase anti-peroxidase soluble complex (PAP, 1:2000, Sigma). eGFP was detected using anti-GFP rabbit polyclonal antibody (1:1000, 1 hr, Abcam) followed by HRP conjugated goat anti-rabbit polyclonal antibody (1:5000, 1 hr, Pierce). TetR was detected using anti-TetR mouse monoclonal antibody (1:1000, 1 hr, Takara). Loading control was detected with mouse monoclonal anti-Ef1α1 (1:15000, 1 hr, Santa Cruz Biotechnologies) and HRP conjugated goat anti-mouse polyclonal antibody (1:2500, 1 hr, Sigma). Membranes were washed thrice with 1X Tris Buffered Saline + Tween 20 (TBST) after each antibody incubation. Chemiluminescent substrate (SuperSignal West Pico Plus, ThermoFisher) was used to detect protein. Band intensities were quantified using ImageJ software (http://imagej.nih.gov/ij/).

#### RNA isolation and northern analysis

5 x 10^7^ cell equivalents were pelleted and washed with PBS for total RNA isolation with TRIreagent (Sigma-Aldrich). RNA was run on a 1.5% agarose/7% formaldehyde gel overnight at 25 V then transferred to a Pall 60208 Biodyne B Membrane as previously described (13). RNA was cross-linked at 1200 J/cm^2^ using a UV Stratalinker 1800 (Stratagene). Specific ^32^P-labelled probes were generated as described previously (14, 19) using PCR products for CymR, TetR, VanR, and POLIB (Table S1) using the Random Primers DNA labeling system (Invitrogen). Tubulin probes were generated from a 650 bp gel extracted product from pZJM (21). Hybridization and wash conditions were described previously (21). Specific mRNAs were detected and quantified using a Typhoon 9500 Phosphorimager (GE Healthcare) and normalized to the tubulin signal.

#### IF microscopy

Cells were harvested at 1000 *x g*, washed once and resuspended in 1X PBS then adhered to poly-L-lysine coated slides (5 min). Cells were fixed with 3 % paraformaldehyde (5 min), washed twice with PBS + 0.1 M Glycine, then permeabilized with 0.05 % Triton X-100 (5 min). Cells were washed thrice with 1x PBS. POLIBWT was detected with rabbit polyclonal anti-protein A (1:1000, 1 hr, Sigma) followed by secondary antibody Alexa Fluor 594 goat anti-rabbit (1:250, 1 hr, ThermoFisher). eGFP was detected using rabbit polyclonal anti-GFP antibody (1:500, 1 hr, Abcam) followed by secondary antibody Alexa Fluor 488 goat anti-rabbit (1:250, 1 hr, ThermoFisher). DNA was stained with 1 µg/ml 4′-6′-diamidino-2-phenylindole (DAPI) and slides were mounted with Vectashield (Vector Laboratories). Images were acquired using an inverted Nikon Ti2-E wide-field fluorescence microscope with a Nikon Plan Apo _λ_ 60x 1.45 numerical aperture objective lens. Z stacks with a step size of 0.2 µm were deconvoluted using the Landweber algorithm within NIS Elements, a closed-source analysis tool developed by Nikon Instruments.

## RESULTS

### Design and Establishing a Triple Inducer System

In our work to expand the molecular toolbox for functional genomics in *Trypanosoma brucei*, we transitioned from using the well establish single inducer Tet-ON system (22) to a dual inducer system for robust RNAi complementation studies. Dual induction relies on the procyclic cell line SMUMA that constitutively expresses Tet and Van repressors as well as T7 RNA polymerase (19). We identified an opportunity to provide additional flexibility by incorporating the previously reported cumate (Cym) inducible components (18, 23) that could allow for the independent regulation of three genetic elements in a single cell line. In this triple inducer system, RNAi complementation would still operate using the Van and Tet inducer elements, while the Cym inducer elements would allow for tunable regulation of additional genes.

Initially, we set out to design a vector that housed both the cumate repressor (CymR) and a cumate (Cym) inducible eGFP reporter under a single drug selectable marker to integrate into the 177 bp locus (Supp Fig1A). We transfected pCuROeGFP into the dual inducer parental cell line SMUMA. Of 10 clonal cell lines, only 4 displayed varied cumate inducible eGFP expression following 48 hr of 25 µg/mL Cym induction with detectable eGFP expression in uninduced controls indicating the system was leaky (Supp Fig1B).

Our second attempt to create a triple inducer cell line relied upon separating the elements onto 2 vectors similar to previous work from the He lab (18). The CymR gene is housed within pCymRHyg to integrate into the α/β tubulin array and the expression vector, pCuO-eGFP, contains 3 cumate operators upstream of an eGFP reporter gene to integrate into the 177 bp repeat region (Fig.1A). SMUMA was transfected with pCymRHyg to create a basal, triple repressor parental cell line called PHITER (<u>P</u>uromycin <u>H</u>ygromycin Inducible <u>T</u>riad of <u>E</u>xogenous <u>R</u>epressors), and 10 clonal cell lines were generated. Several PHITER clones were subsequently transfected with pCuO-eGFP to characterize cumate inducible expression using eGFP as a reporter. Following induction with 25 µg/mL Cym for 48 hr, four eGFP^PHITER^ populations showed varying levels of eGFP protein expression while no expression was detected in the uninduced cells (Fig. 1B). Longer exposures (90 sec) showed no expression of eGFP in the uninduced controls (Supp Fig1C). These results demonstrate that separating the Cym elements onto 2 vectors eliminated the problem of leakiness in the uninduced controls.

**FIG 1.**
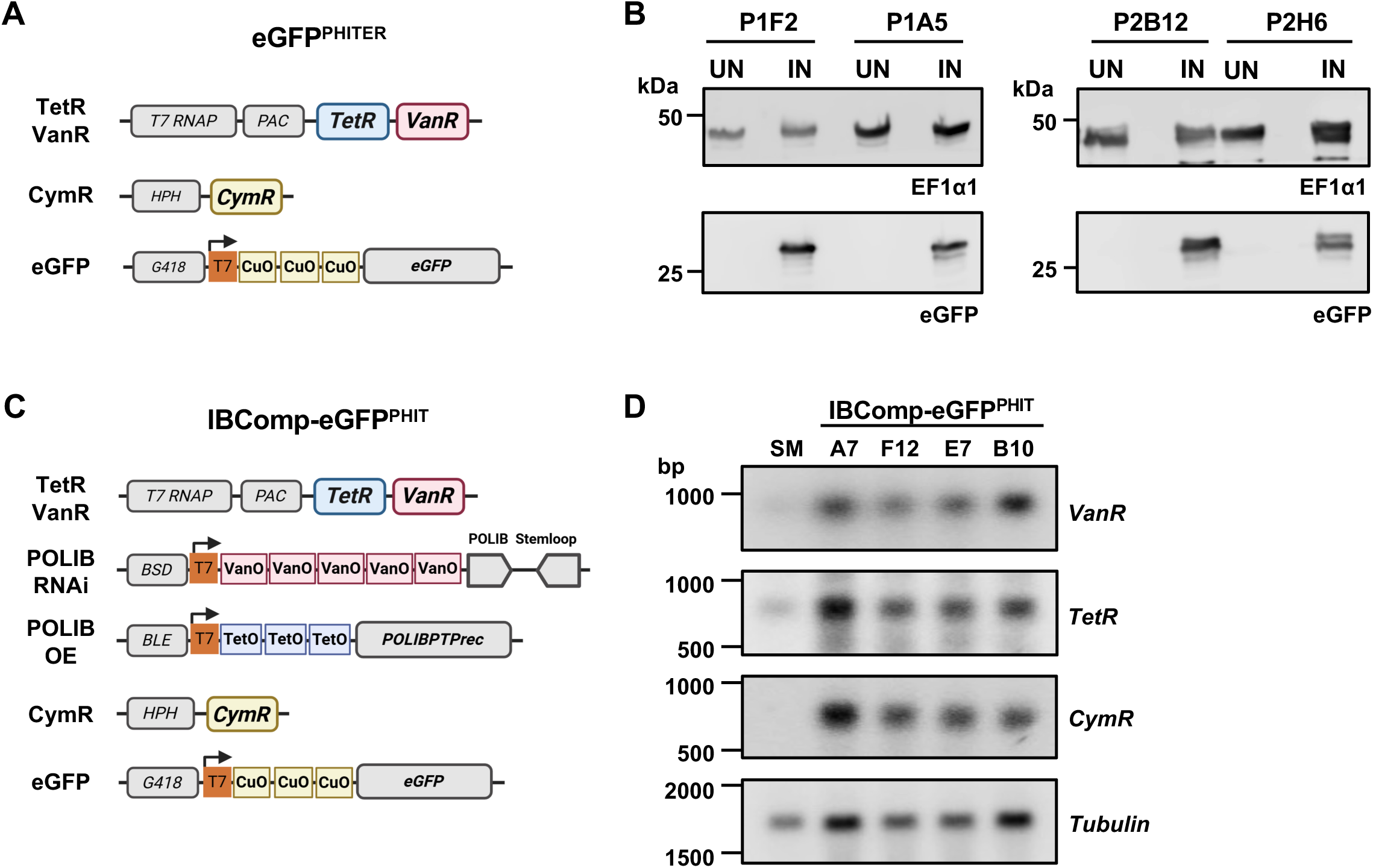
Triple Inducer Systems. **(A)** Diagram of plasmids integrated into the eGFP^PHITER^ cell line. Tetracycline repressor (TetR), vanillic acid repressor (VanR), T7 RNA polymerase (T7RNAP), puromycin-N-acetyl-transferase (PAC), cumate repressor (CymR), hygromycin B phosphotransferase (HPH), neomycin phosphotransferase II (G418), and cumate operator (CuO). **(B)** Western blot detection of eGFP and loading control EF1α1 following a 48 hr induction of eGFP^PHITER^ clones with 25 μg/mL Cym. 2 x 10^6^ cell equivalents were loaded per lane. **(C)** Diagram of plasmids integrated into the IBComp-eGFP^PHIT^ cell line. Blastacidin S-deaminase (BSD), vanillic acid operator (VanO), tetracycline operators (TetO), bleomycin resistance gene. Other annotation as described in A. **(D)** Northern blot of total RNA from IBComp-eGFP^PHIT^ clonal cell lines. SMUMA, parental cell line as a control. Membrane was probed for individual repressor mRNAs, stripped and then reprobed. *Tubulin*, loading control.

To demonstrate independent control of all three inducers, we adapted the previously characterized RNAi complementation cell line, IBComp^VaT^ (19), to work in parallel with the Cym inducible elements. IBComp^VaT^ was transfected with pCymRHyg and pCuO-eGFP to create IBComp-eGFP^PHIT^ thus allowing Cym inducible eGFP expression in addition to Van inducible POLIB RNAi and Tet inducible POLIBWT overexpression within a single cell line (Fig. 1C). Both the pCymRHyg and pJ1173 integration events occur within the α/β tubulin repeat region. Therefore, we performed northern blot analyses to confirm expression of each repressor mRNA in the 4 clones using SMUMA cells as a control (Fig. 1D). IBComp-eGFP^PHIT^ clone P1F12 was selected as the cell line to analyze proof-of concept for triple induction based on repressor expression, and similar POLIB RNAi complementation results as previously published (19).

### POLIB Triple Inducer System: Optimization of Cumate Inducible eGFP

To determine if the triple inducer cell line IBComp-eGFP^PHIT^ was able to express eGFP across a range of Cym concentrations (0.5 – 10 μg/mL) were added to the cells for 48 hr, cell pellets were collected and then analyzed by western blot analyses. No eGFP expression was observed in uninduced cells even upon a longer exposure of 5 min (Fig. S2A). Expression of eGFP was detected from 1 μg/mL Cym and consistently increased as the Cym concentration increased to 10 μg/mL (Fig. S2A). Cells induced with 5 μg/mL of Cym or greater showed degradation of eGFP (Fig. S2A). We further tested 3 and 5 μg/mL Cym for sustained expression of eGFP over an 8 day induction. While some variation in the amount of eGFP was detected using 5 μg/mL, there was consistent expression of eGFP using 3 μg/mL Cym throughout the 8 day induction (Fig S2B). We also tested whether Cym concentrations impacted fitness of the cells. No significant fitness impact was detected by inducing the cells with a range of Cym concentrations (3, 5, 10 μg/mL) or by expression of eGFP over an 8 day induction (Fig. S2C). Based on these data, 3 μg/mL Cym was used for subsequent inductions.

### Independent Gene Expression in IBComp-eGFP^PHIT^ Cell Line

One advantage of the triple inducer system is that all the single inducer controls can be performed in a single cell line. To determine whether the Van, Tet and Cym inducible systems could operate independently of one another, each inducer was added separately to the IBComp-eGFP^PHIT^ cell line. Cells were cultured for 8 days with the addition of either 3 μg/mL Cym, 4 μg/mL Tet, or 250 μM Van. After 48 hours, ectopic POLIBWT and eGFP expression were assessed by western blotting and endogenous *POLIB* mRNA was quantified via northern blotting. Cells induced with Tet expressed POLIBWT (10.1 fold greater than SE) with no detectable eGFP, while cells induced with Cym expressed eGFP with no detectable expression of POLIBWT (Fig 2A).

**FIG 2.**
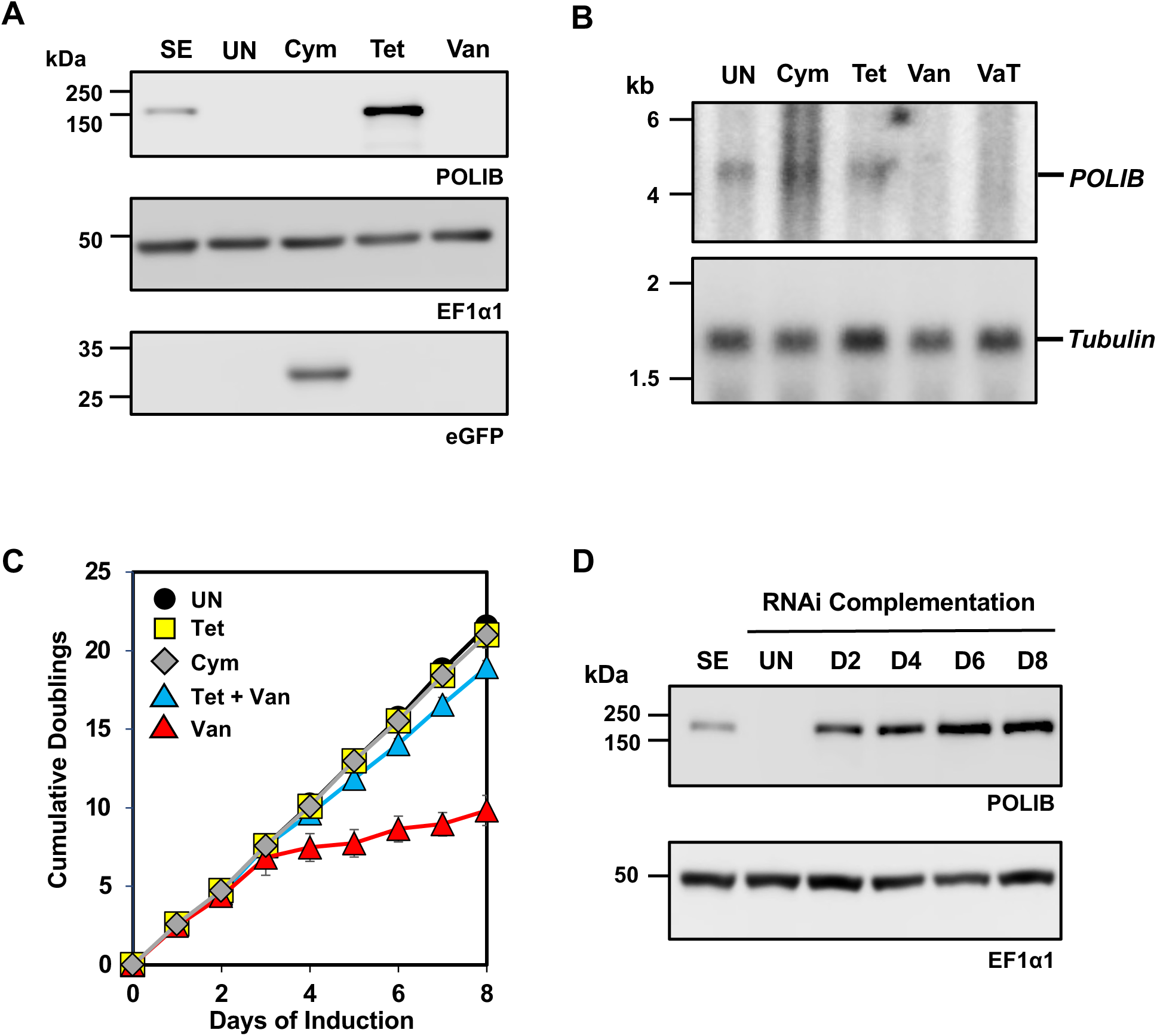
Independent gene expression in IBComp-eGFPPHIT cells. **(A)** Western blot detection of POLIB, eGFP and EF1α1 from IBComp-eGFP^PHIT^ cells independently induced with Tet (4 μg/ml), Van (250 μM) Cym (3 μg/ml) over an 8 day induction. SE, single expressor cell line representing endogenous POLIB protein (4 x 10^6^ cell equivalents). 2 x 10^6^ cell equivalents were loaded per lane. **(B)** Northern blot of total RNA from IBComp-eGFP^PHIT^ cells. U, uninduced; Cym, induced with 3 μg/ml Cym for 48 hr; Tet, induced with 4 μg/ml Tet for 48 hr; Van, induced with 250 μM Van for 48 hr; VaT, induced with Van and Tet for 48 hr. Top, probing of *POLIB* mRNA. Bottom, probing of *Tubulin* mRNA as a loading control. **(C)** IBComp-eGFP^PHIT^ was grown in the absence of presence of Tet (4 μg/ml), Van (250 μM) Cym (3 μg/ml), and Tet and Van combined. Error bars represent ± s.d. of the mean from 3 biological replicates. **(D)** Western blot detection of IBWT and loading control EF1α1 from IBComp-eGFP^PHIT^ grown in 250 μM Van and 4 μg/ml Tet for 8 days. SE, single expressor cell line control. 2 x 10^6^ cell equivalents were loaded per lane.

Induction with Van did not lead to expression of either POLIBWT or eGFP while endogenous *POLIB* mRNA was depleted by 98% within 48 hr (Fig. 2B). Throughout the 8 day induction only the addition of Van led to a significant loss of fitness (LOF) with cell doublings reaching only 9.8 compared to 21.5 for uninduced cells, a hallmark associated *POLIB* RNAi (12, 14, 19). Cells induced with Tet and Cym reached 20.8 and 21 doublings, respectively (Fig. 2C). These data confirm that the three inducers operate independently and did not lead to promiscuous expression.

To determine if POLIB RNAi complementation was still functioning in the triple induction system, IBComp-eGFP^PHIT^ cells were simultaneously induced with Van and Tet for 8 days. Consistent expression of POLIBWT was confirmed with a 7 fold increase compared to the single expressor allelically tagged endogenous control (SE) (Fig. 2D). Similar to RNAi alone, induction with Van and Tet resulted in a 99% knockdown of endogenous *POLIB* mRNA after 48 hr of induction (Fig. 2B). Lastly, there was a near complete rescue of the RNAi LOF during POLIB complementation with cells reaching 18.9 doublings by day 8 phenocopying results for the dual inducer system (19) (Fig. 2D).

### Selective and Concurrent Gene Expression in IBComp-eGFP^PHIT^ Cell Line

To assess whether Cym induction interfered with Van and/or Tet inductions we next evaluated whether these inducers could operate simultaneously in IBComp-eGFP^PHIT^ cells. The following combinations of inducers were used across 8 days of induction; Van/Cym (VC), Tet/Cym (TC), and Van/Tet/Cym (VTC) and assessed for POLIB mRNA knockdown, POLIBWT expression, and eGFP expression after 48 hr. TC induction did not have a significant impact on fitness (21.2 doublings) compared to uninduced cells (21.6 doublings) and led to the highest amount of POLIBWT and eGFP expression with no impact on endogenous *POLIB* mRNA (Fig. 3A, B, C). VC induction resulted in a phenocopy of RNAi alone (Van) with the characteristic LOF (9.5 doublings), a 94% reduction in *POLIB* mRNA and expression of eGFP (Fig. 3A, B, C). The full triple induction (VTC) also led to a 97% reduction in *POLIB* mRNA, a similar near complete rescue of the RNAi LOF and eGFP expression (Fig. 3A, B, C).

**FIG 3.**
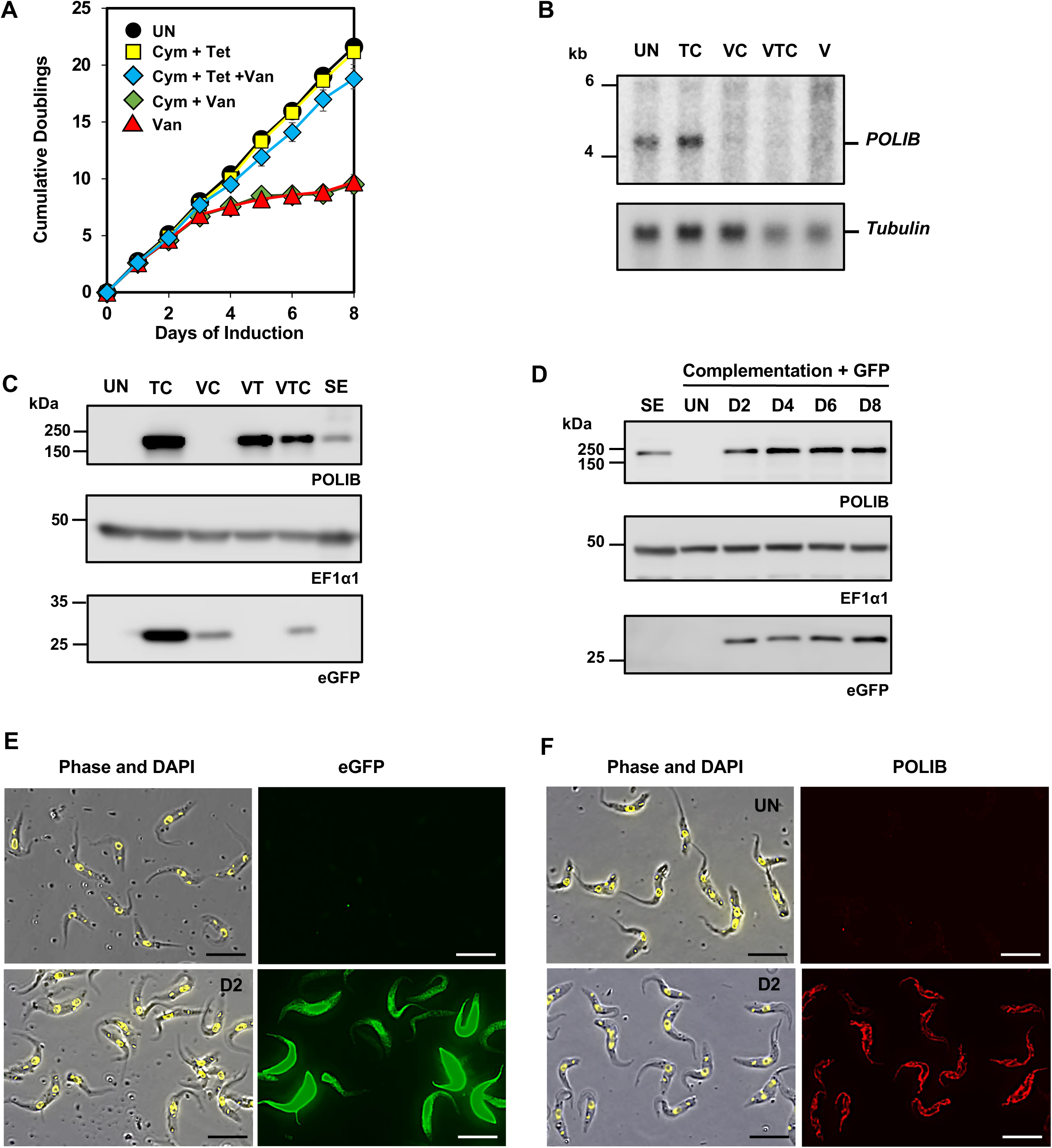
Concurrent Gene Expression in IBComp-eGFP^PHIT^ cells. **(A)** IBComp-eGFP^PHIT^ was grown in the absence or presence of Van (250 μM), or inducer combinations; Cym (3 μg/ml) and Tet (4 μg/ml), Cym and Van, and Cym, Tet, and Van for 8 days. Error bars represent ± s.d. of the mean from 3 biological replicates. **(B)** Northern blot of total RNA from IBComp-eGFP^PHIT^ cultured for 48 hours in the presence of inducer combinations. U, uninduced; TC, Tet and Cym; VC, Van and Cym; VTC, Van, Tet and Cym; V, Van. Top, probing of *POLIB* mRNA. Bottom, probing of *Tubulin* mRNA as a loading control. **(C)** Western blot of POLIB, eGFP and EF1α1 loading control from IBComp-eGFP^PHIT^ cells concurrently induced with Cym, Tet, and Van for 8 days. SE, single expressor cell line control. 2 x 10^6^ cell equivalents were loaded per lane. **(D)** Western blot of eGFP, POLIB and EF1α1 loading control from IBComp-eGFP^PHIT^ cells concurrently induced with Cym, Tet, and Van for 8 days. SE, single expressor cell line control. 2 x 10^6^ cell equivalents were loaded per lane. **(E)** Representative images of eGFP expression for IBComp-eGFP^PHIT^ in the presence or absence of Cym+Tet+Van for 48 hours. DAPI staining (yellow); eGFP expression (green). Size bar, 10 μm. **(F)** Representative images of POLIB IBComp-eGFP^PHIT^ in the presence or absence of Cym+Tet+Van for 48 hours. DAPI staining (yellow); anti-protein A (red). Size bar, 10 μm.

Compared to Tet alone, TC induction led to a 12.3 fold increase in POLIBWT, whereas VT induction led to a slight decrease in POLIBWT (7 fold) that did not significantly impact *POLIB* RNAi complementation (Fig. 2C, D, Fig. 3C). The triple induction with VTC resulted in a 6.5 fold above endogenous POLIB levels that still allowed for complementation. For VC, eGFP decreased 3.2 fold compared to Cym alone. With a comparable decrease of 3.6 fold for the triple VTC inductions (Fig 3C). Although the addition of Van seemed to impact the total amount of eGFP or POLIBWT being produced, the simultaneous induction of three inducers still led to a near complete rescue of the *POLIB* RNAi LOF with 18.8 doublings with consistent and sustained expression of eGFP and POLIBWT (Fig 3A, D).

Lastly, fluorescence microscopy revealed that eGFP and POLIBWT expression were homogenous in the cell population with POLIBWT localizing throughout the mitochondrion and concentrating near the kDNA while eGFP was cytoplasmic with some fluctuation in the total amount of protein being produced (Fig. 3E, F). These data indicate that Cym induction did not significantly interfere with *POLIB* RNAi or overexpression of POLIBWT and highlight that Van induction might lead to decreased expression levels for genes under the control of Cym and Tet induction.

### Temporal Expression of eGFP during RNAi Complementation

Another advantage of the triple induction system would be to control when a gene of interest is expressed while performing RNAi complementation. Therefore, we tested whether eGFP expression could be controlled temporally during Cym induction and *POLIB* RNAi complementation. Following 48 hr of Cym induction, removal of Cym resulted in a decrease in eGFP expression within 24 hr of removing Cym with some residual eGFP expression detected the following day (Fig. 4A). Similarly, removal of Cym during RNAi complementation resulted in more rapid reduction of eGFP expression within 24 hr after removing Cym (Fig. 4B). Collectively these data confirm the specificity of three independent inducers, ability to tune the amount of Cym and to control when Cym induction occurs. Together the results establish proof of principle for a triple inducer system in procyclic *T. brucei* that can be used for elegant molecular genetic studies during RNAi complementation.

**FIG 4.**
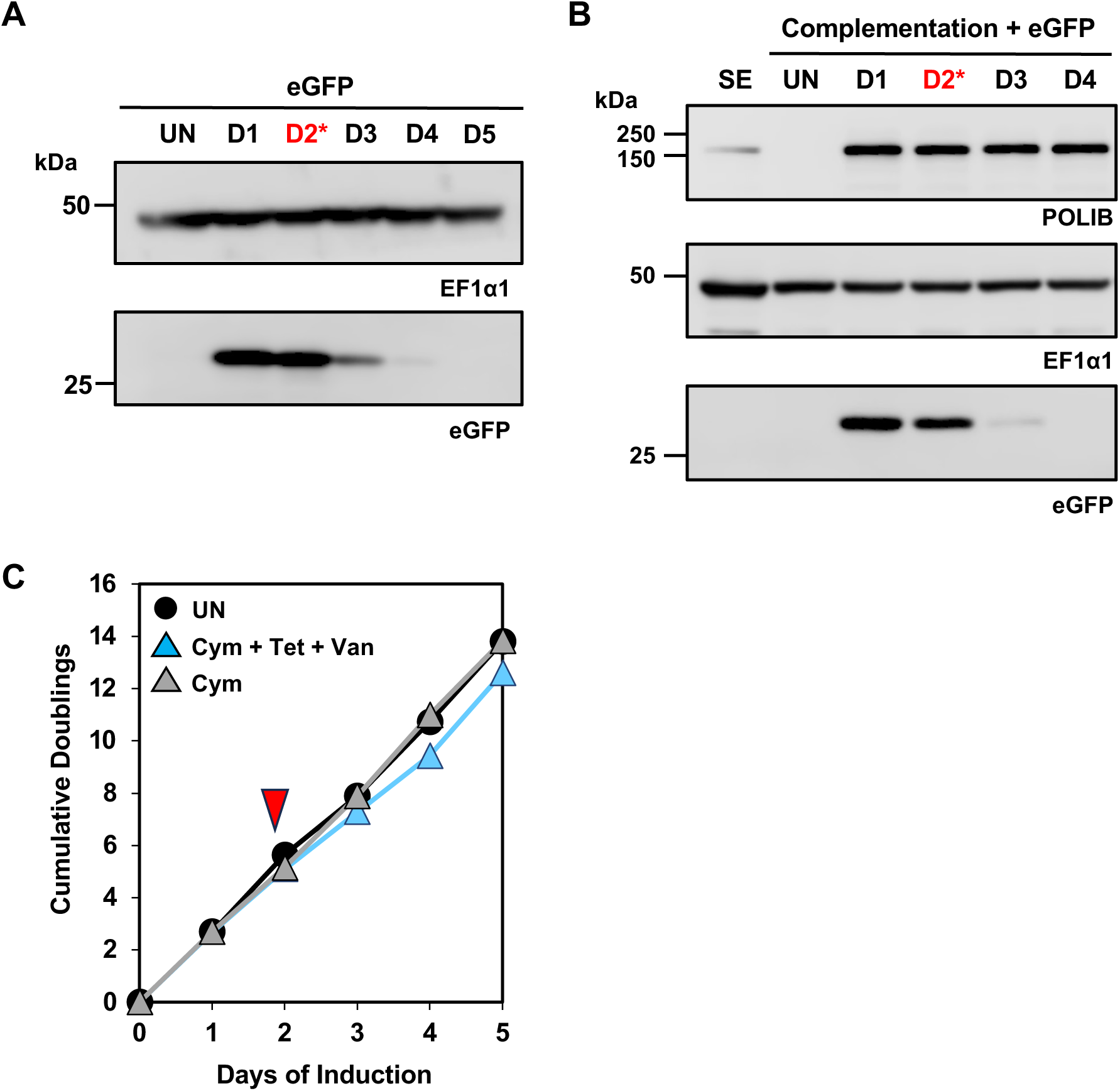
Temporal Expression of eGFP in the Triple Inducer System. **(A)** Western blot detection of eGFP and EF1α1 loading control from IBComp-eGFP^PHIT^ cells induced with Cym (3 μg/ml), for 2 days then in the absence of Cym for subsequent days. D2*, sample taken from cell before Cym inducer was removed. 2 x 10^6^ cell equivalents were loaded per lane. **(B)** Western blot detection of eGFP, POLIB and EF1α1 loading control from IBComp-eGFP^PHIT^ cells induced with Van, Tet and Cym for 2 days then in the absence of Cym for subsequent days. D2*, sample taken from cell before Cym inducer was removed. SE, single expressor cell line control. 2 x 10^6^ cell equivalents were loaded per lane. **(C)** IBComp-eGFP^PHIT^ was grown in the absence or presence of Cym (3 μg/ml) or all inducers for 5 days. Red triangle, Cym inducer was removed and grown in absence of Cym. Graphs represent averaged data from two biological replicates.

## Discussion

The use of inducible expression for reverse genetics and functional genomics screens has been critical for studying gene function in the model kinetoplastid *T. brucei.* Advances include several RNAi library screens, and an overexpression gain-of-function screen (16, 24). However, there are limitations to the single inducer Tet-On system that have prohibited powerful multiplexed screens (25). Here, we have established a highly tractable, triple inducer system capable of independently or simultaneously inducing the expression of three ectopic genes within one cell line. A triple inducer cell line allows for more flexibility to now perform sophisticated genetic screens, and mechanistic studies that could never have been achieved with the single inducer Tet-On system or the recently reported dual inducer system for RNAi complementation (19).

We first attempted to engineer a parental cell line for triple induction that only required one additional drug marker to allow for more flexibility in the design of downstream experiments. However, the pCuRO-eGFP construct that contained both the CymR gene, and a cumate inducible reporter gene (eGFP) only produced clones that displayed leaky eGFP expression (Fig S1B). At this time, we are not certain why this arrangement and integration into the 177 bp repeat region resulted in uncontrolled eGFP expression (Fig S1A B).

The basis for the triple inducer system is a parental cell line called PHITER (derived from SMUMA) which separated the Cym elements onto two vectors for integration at separate genomic loci. By capitalizing on the previously reported inducible elements using Tet, Van and Cym to control gene expression in *T. brucei* (26, 17, 18), we demonstrated constitutive expression of all the repressor elements CymR, VanR, and TetR (Fig S3), and characterized cumate inducible eGFP in a single cell line (Fig 1B). Longer exposures of up to 2 minutes confirmed that there was no evidence of leaky expression in the uninduced cells. Clonal PHITER cell lines have doubling times (9-10 hrs) comparable to the parental strain SMUMA and are available to the field. In contrast, the widely used 29-13 cell line for the single inducer Tet-On system grows more slowly (∼12.8 hr), expresses lower levels of TetR, and displays considerable divergence in the minicircle genome (27).

As proof of concept, we evaluated the full potential of the triple inducer system using the previously characterized Van/Tet dual inducer cell line for POLIB RNAi complementation, IBComp^VaT^ (19). Consecutive transfections of pCymRHyg then pCuO-eGFP produced IBComp-eGFP^PHIT^, a cell line that could be used for RNAi and ectopic expression of two genes independently or simultaneously (Fig 1C). Independent induction with Cym, Tet, and Van was demonstrated with no evidence of promiscuous expression (Fig 2A, B). Van induction resulted in *POLIB* mRNA depletion (98-99% decreased) and Tet induction resulted ectopic expression of IBWT (10.1 fold) that closely resembled those in the parental IBComp^VaT^ experiments (Fig 2A, B). In addition to controlling gene expression independently, we also demonstrated that Cym induction of eGFP could be tunable over a range of Cym concentrations (1 - 10 µg/mL) and that there was sustained expression of Cym induced eGFP (Fig S2C, D). Even the highest concentration of Cym do not impact fitness of cells when tested in the parental cell line SMUMA (Fig S2C). Importantly, Van + Tet dual induction for RNAi complementation led to a near complete rescue of the POLIB RNAi defect and sufficient overexpression of ectopic protein (Fig 2C, D).

We also confirmed concurrent gene expression with various dual and finally the full triple inducer conditions. Combining Van + Cym still resulted in the characteristic 98% POLIB mRNA depletion and LOF, and Cym +Tet resulted in robust expression of both eGFP and IBWT (Fig 3). Induction with all three inducers allowed for sustained expression of ectopic POLIB (Fig 3D), expression of eGFP, knockdown of endogenous POLIB (Fig 3C), and a near complete rescue of the RNAi fitness defect (Fig 3A). These results demonstrated that all inducer combinations caused the expected expression output.

Another important aspect of the triple inducible system is the ability to control timing of independent inductions for more dynamic experimental applications. Control of RNAi (removal of the Van inducer) was previously demonstrated by removing the Van inducer to restore the expression of chromosomally epitope tagged POLIB (19). Here we additionally demonstrate the temporal control of Cym inductions by removing the Cym inducer after 48 hrs of eGFP expression. In single or the full triple induction conditions, removal of Cym inducer resulted a rapid decline in eGFP expression within 24 hrs. During the triple induction, POLIBWT ectopic expression remained consistent throughout the induction demonstrating that cumate inducible protein could be temporally expressed without impacting concurrent RNAi complementation (Fig 4C).

When designing the dual and triple inducer systems, relative fold increase in expression levels were taken into consideration. To maximize overexpression of an ectopic protein during RNAi complementation, we placed POLIBWT on a Tet expression vector that had previously reported to give a greater fold increase compared to the Van expression vector (250 fold vs 18 fold increase in fluorescence intensity (17). The POLIB RNAi stemloop construct described previously (28) was then placed on a Van expression vector and efficient RNAi knockdown using this system was previously reported (17, 19). Even though fold expression using Van induction was substantially lower, the long dsRNA constructs used in *T. brucei* provide ample siRNAs for efficient knockdown. The cumate inducible system was then used for expression of a third gene independent of RNAi complementation due to the previously demonstrated ability to tune the amount of protein and temporally control expression (18). We have not tried other design combinations of the triple inducible system.

However, the Cym, Tet, and Van inducible elements each have limitations that should be considered during experimental design. Tet is a common contaminant in FBS used to culture parasites possibly leading to low levels of background protein expression.

Previously, 250 µM Van was reported to cause a slight fitness defect in *T. brucei* (19) and has been demonstrated to exhibit dose-dependent anti-trypanosomal activity in *T. congolensi* with a reported IC50 of 768 µg/ml. (29). Lastly, Cym and Tet inducible protein expression is reduced in concurrent inductions with Van (Fig 3D), thus the concentration of Cym used in combination with Van may need to be titrated to circumvent any observed reduction in protein expression.

Our study provides proof of concept for the first dynamic triple control system that allows for robust RNAi complementation and control of a third element based on Tet, Van and Cym inducible gene expression, and expands the toolbox available for *T. brucei* functional genomics. Similar to the dual induction system, the triple system allows for independent and tunable gene expression to control multiple processes. Additionally, the three elements can also be temporally modulated for simultaneous or more elegant staggered (or contemporaneous) experimental designs such as overexpressing proteins that specifically modulate kDNA stability prior to performing RNAi complementation.

---

**Supplemental Fig 1.** Optimization of the Cumate Inducer System. (A) Diagram of plasmids integrated into the CuRO-eGFP cell line. Cumate repressor (CymR), (HPH), neomycin phosphotransferase II (G418), cumate operator (CuO), tetracycline repressor (TetR), vanillic acid repressor (VanR), T7 RNA polymerase (T7RNAP), puromycin-N-acetyl-transferase (PAC). **(B)** Western blot detection of eGFP and EF1α1 loading control from CuRO-eGFP clonal cell lines. 2 x 10^6^ cell equivalents were loaded per lane. **(C)** Western blot detection of eGFP and EF1α1 loading control from eGFP^PHITER^ clonal cell lines. 2 x 10^6^ cell equivalents were loaded per lane. Image represents a longer exposure (90 seconds) of the data presented in Figure 1B.

**Supplemental Fig 2.** Cumate Inducible eGFP Expression in IBComp-eGFP^PHIT^. **(A)** Western blot detection of eGFP and EF1α1 loading control from IBComp-eGFP^PHIT^ cells induced with varying concentration of Cym for 48 hr. 2 x 10^6^ cell equivalents were loaded per lane**. (B)** Western blot detection of eGFP and EF1α1 loading control from IBComp-eGFP^PHIT^ cells induced with varying concentration of Cym for 8 days. **(C)** IBComp-eGFP^PHIT^ was grown in the absence of presence of varying concentrations of Cym. Graphs represent averaged data from two biological replicates.

**Supplemental Fig 3. Characterization of PHITER clonal cell lines. (A)** Western blot detection of TetR and EF1α1 loading control from PHITER clonal cell lines. 2 x 10^6^ cell equivalents were loaded per lane**. (B)** Northern blot of total RNA from PHITER clonal cell lines. Left, probing for *CymR* mRNA. Right, probing for *VanR* mRNA. Bottom for each, rRNA as a loading control.

