## supplemental information for "A Trypanosome Trifecta: an independently tunable Triple Inducer System for genetic studies in *Trypanosoma brucei*"

**Supplemental Fig 1** : Optimization of the Cumate Inducer System

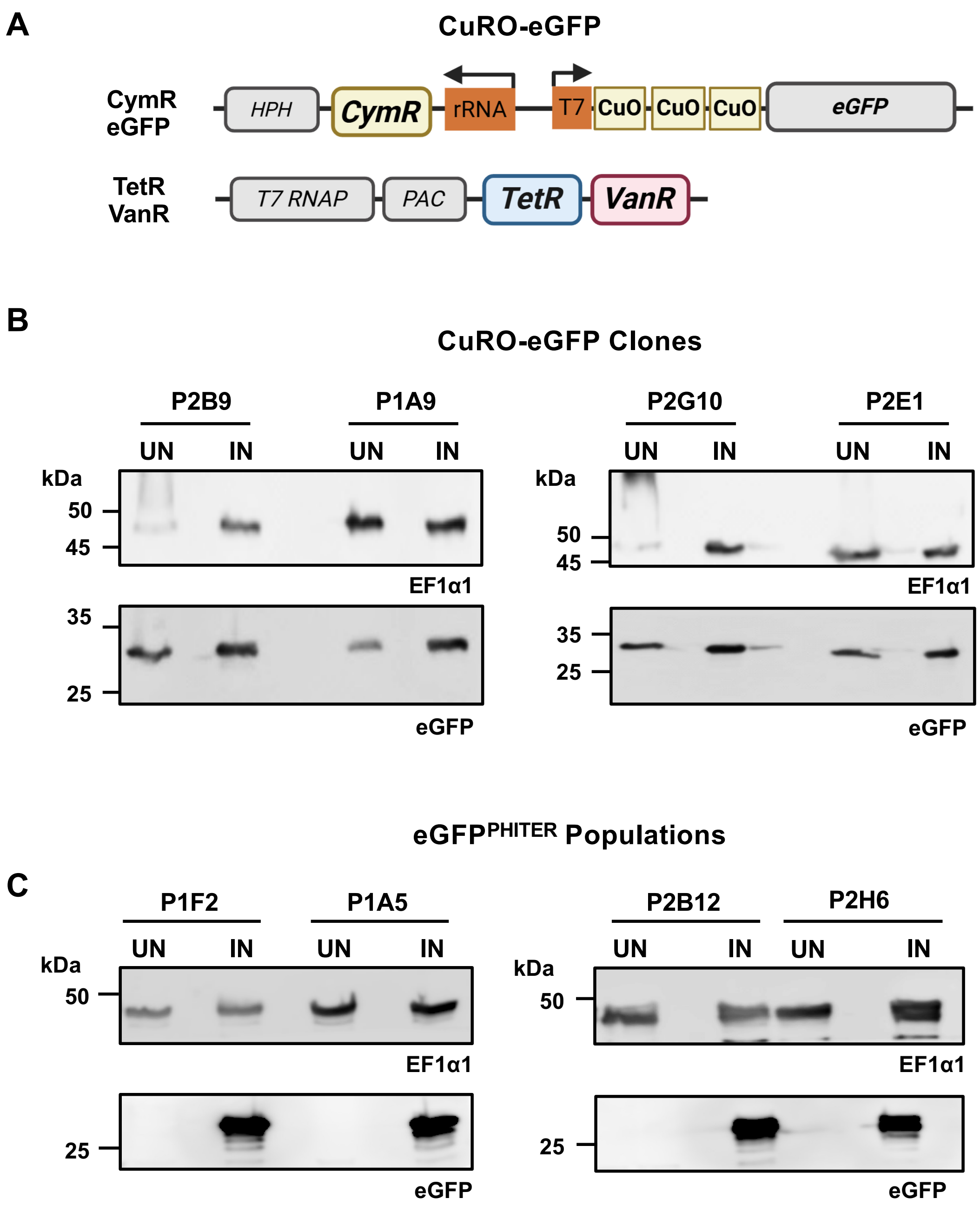

**Supplemental Fig 2** – Cumate Inducible eGFP Expression in IBComp-eGFP$^{PHIT}$

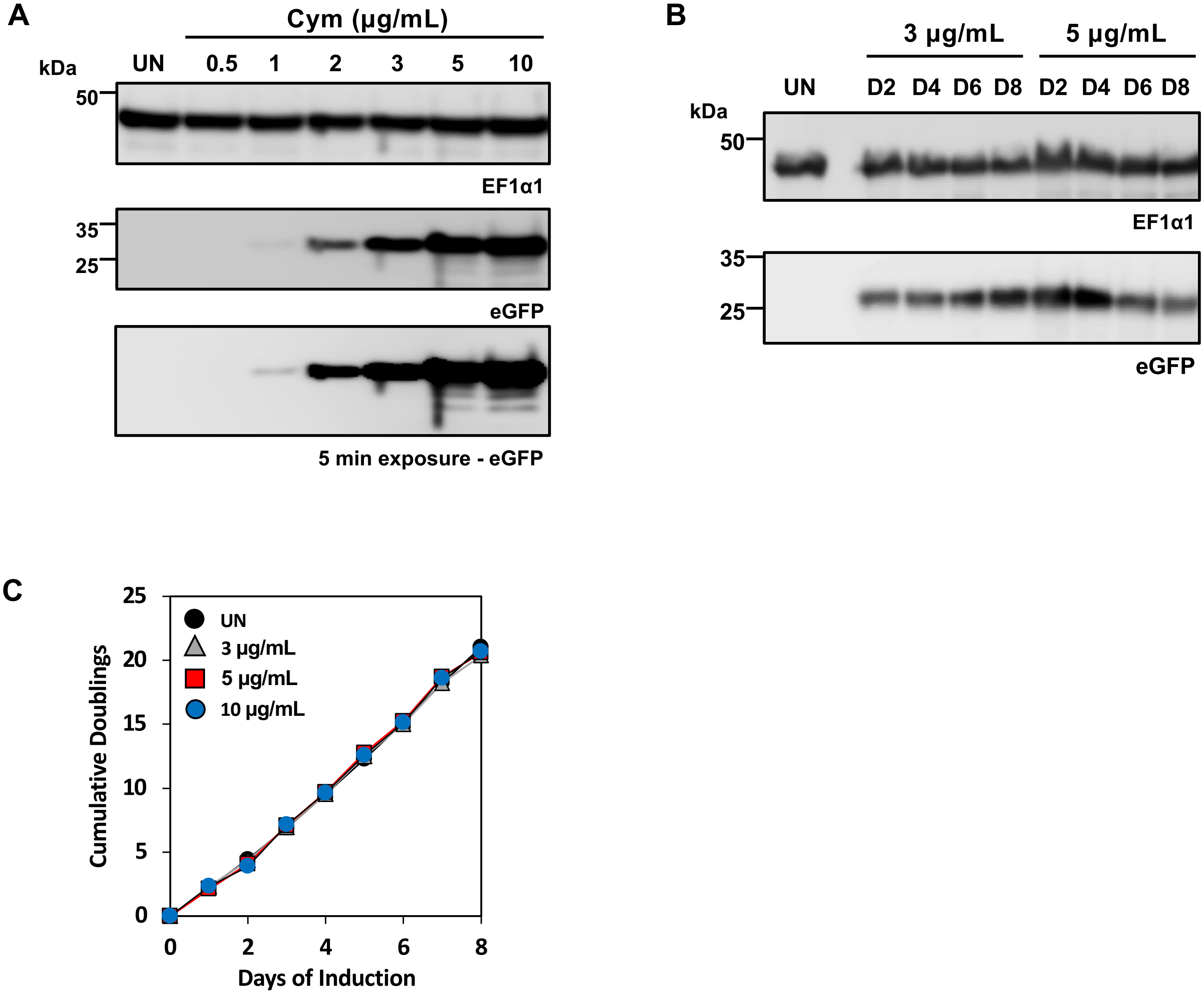

**Supplemental Fig 3** : Characterization of PHITER clonal cell lines

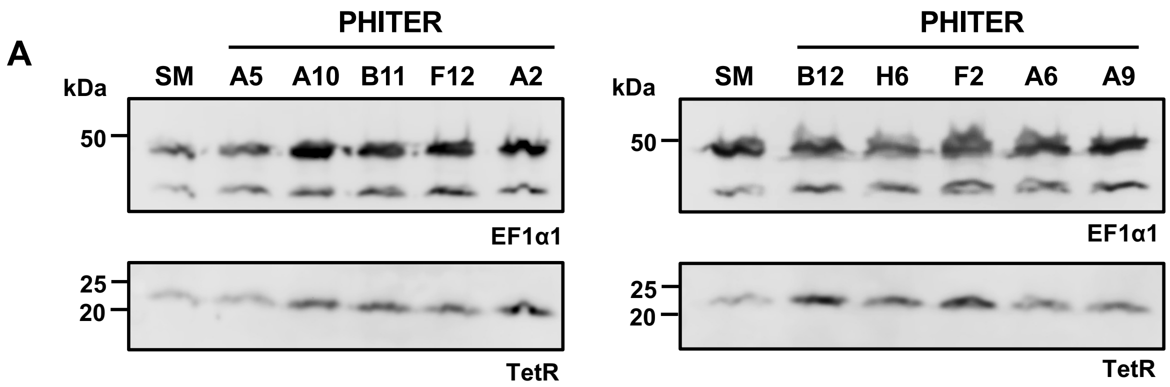


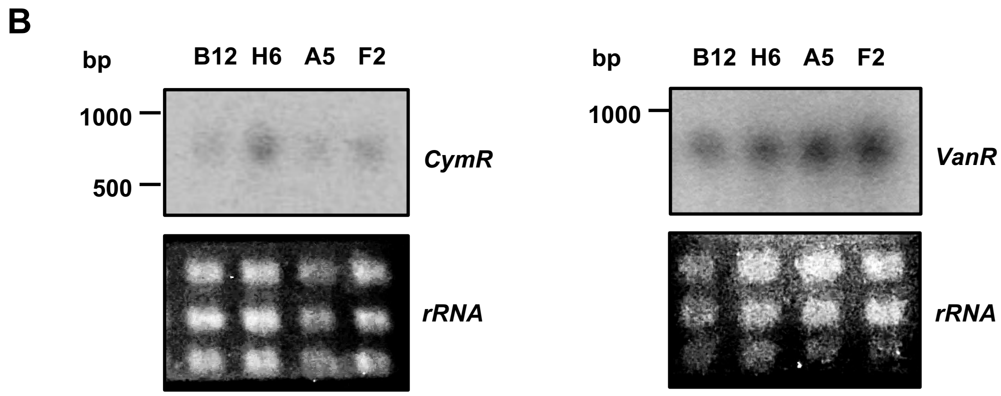

**Table S1: Primers used in this study.**

| Gene | Purpose | Primer Name | Primer Sequence (5' – 3') |
|---|---|---|---|
| Hygromycin-B phosphotransferas WVW91683.1 | Gibson Cloning pSmOxNUSHyg | UM147 | AGC AAT AAA GCA TCA GAA TTA TGA AAA AGC CTG AAC TCA CCG C |
| | | UM149 | CAC TTA AGC GCA GCG CCA TGC TAT TCC TTT GCC CTC GGA CGA |
| Not applicable | Gibson Cloning pCuRO-eGFP | UM159 | CGG TGT TAG GAT CTC CGA GGT TGT GGC CGC GCA TCC TAG G |
| | | UM160 | AAG TAG CGC TTA CGG CGT ACC GCG TTC GCG TAA GGA TCC C |
| Not applicable | Gibson Cloning pCymRHyg | UM192 | AAA ATA GTT CAA ACG AAT TAG GCA ACC TGA ACC TTC GC |
| | | UM193 | GGA AAT GCC CCG TCC GCG TGC GCC ATG CTA TTC CTT TGC |
| Neomycin phosphotransferase CAD21776 | Gibson Cloning pCuO-eGFP | UM184 | CGG TGT TAG GAT CTC CGA GGT CAG AAG AAC TCG TCA AGA AG |
| | | UM185 | GCC ATA AAA TAA GCT ATC ACA TGA TTG AAC AAG ATG GAT TGC ACG |
| POLIB Tb927.11.4690 | Northern blot | MK157 | CAT TCA CAG GGG TTG AAG TCA TCG C |
| | | MK159 | ACG TTC CAC CCT ACT GTA CAC TAC G |
| CymR O33453.1 | Northern blot | UM212 | GCG TGA TTG CTT AGT AAG TTG GTG |
| | | UM213 | TAC GAA TCA GTG CCC AAG TGG AGC |
| VanR WP_010920250.1 | Northern blot | UM125 | GCC AGG TCA GCG TGT GAT GAT G |
| | | UM126 | TGC TCG TAC TCT GCT GGA AGA TCC |
| TetR XNS37014.1 | Northern blot | UM236 | GCCTGGACAAGAGCAAGGTCATTAAC |
| | | UM237 | AACCCGACTCGCACTTGAGCTGCTTC |

### Table S2: Cell lines used in this study

| Name | Description | Drug Selection | Reference |
|---|---|---|---|
| SMUMA (Dual Inducer) | Lister 427<br>T7RNAP + TetR + VanR | Puro | Armstrong et al. 2025 |
| IBPTP/IBKO | Single Expressor POLIB-PTP | G418, Puro | Armstrong et al. 2025 |
| IBComp$^{VaT}$ | SMUMA<br>POLIB RNAi + POLIB-PTP | Puro, Bsd, Phleo | Armstrong et al. 2025 |
| PHITER (Triple Inducer) | SMUMA<br>CymR | Puro, Hyg | This Study |
| eGFP$^{PHITER}$ | PHITER<br>eGFP | Puro, Hyg, G418 | This Study |
| IBComp$^{PHIT}$ | IBComp$^{VaT}$<br>CymR | Puro, Bsd, Phleo, Hyg | This Study |
| IBCompeGFP$^{PHIT}$ | IBComp$^{PHIT}$<br>eGFP | Puro, Bsd, Phleo, Hyg, G418 | This Study |
